# Virus-induced upregulation of mitochondrial metabolism modulates cytosolic redox balance and defense responses

**DOI:** 10.1101/2025.07.05.663277

**Authors:** Liang-Yu Hou, Chih-Hang Wu, Na-Sheng Lin

**Author notes:** Corresponding author: Dr. Na-Sheng Lin, Address: Institute of Plant and Microbial Biology, Academia Sinica, 115201, Taipei, Taiwan. N. -S.L. and L. -Y.H. conceived the project; N. -S.L. and L. -Y.H. designed the research; L. -Y.H. performed the experiments; N. -S.L. and L. -Y.H. analyzed the data; N. -S.L. and C. -H.W. supervised the experiments; N. -S.L., C. -H.W. and L. -Y.H. wrote the manuscript. This work was supported by Academia Sinica Investigator Award (AS-IA-108-L05) and Academia Sinica Postdoctoral Scholar Program. The author responsible for distribution of materials integral to the findings presented in this article in accordance with the policy described in the Instructions for Authors (www.plantphysiol.org) is Na-Sheng Lin.

## Abstract

Plants possess a remarkable capacity to reprogram their metabolism in response to pathogen attacks. However, the mechanisms by which metabolic reprogramming modulates defense signaling remain poorly understood. In this study, we leverage a multifaceted omics approach to investigate the metabolic shifts induced by the Bamboo mosaic virus (BaMV), a positive-sense single-stranded RNA virus that depends on host factors from multiple organelles for its replication. Metabolic profiling revealed an accumulation of hexose phosphates and Krebs cycle intermediates in *Nicotiana benthamiana* plants following BaMV infection. Fluxomic analysis uncovered an orchestrated redirection of metabolic flux toward glycolysis and the Krebs cycle during infection. Proteomic data further highlighted a concerted upregulation of mitochondrial enzymes, with three mitochondrial proteins showing markedly increased accumulation in BaMV-infected tissues. These integrated omics results suggest that BaMV infection triggers a metabolic shift toward energy-generating pathways. Notably, functional analysis revealed that silencing mitochondrial NAD^+^-dependent malic enzyme 1 significantly enhanced BaMV accumulation, accompanied by alterations in cytoplasmic NADH-to-NAD^+^ ratio and changes in the landscape of defense gene expression. Collectively, our findings underscore the pivotal role of mitochondrial metabolism in governing cytoplasmic redox balance, finely tuning defense responses to viral infection.

**One sentence summary:** BaMV infection enhances mitochondrial metabolism to regulate cytoplasmic redox balance and promote antiviral defense.

## Introduction

In the intricate arms race between plants and viruses, both organisms have evolved sophisticated strategies to gain an advantage. Viruses, as obligate intracellular parasites, hijack host cellular machinery to ensure their replication, movement, and survival. In turn, plants have evolved multilayered immune responses, ranging from RNA silencing to hormone-mediated defense signaling, aimed at recognizing and counteracting viral invasion (Alazem et al., 2014; Alazem and Lin, 2017; Cheng and Wang, 2017; Wu et al., 2019; Alazem and Lin, 2020; Fontes et al., 2021; Ge et al., 2024). This dynamic interplay shapes the outcome of infection and often centers around the host’s ability to remodel cellular physiology in response to viral cues.

Metabolism, a dynamic network of biochemical reactions, lies at the heart of cellular function and survival. Initiating such metabolic acclimation is critical for plants to adapt to biotic stress, including viral infections (Rojas et al., 2014; Llave, 2016; Kanwar and Jha, 2019; Xiao and Loscalzo, 2020; Xu et al., 2022; Jiang et al., 2025). As the frontline of this molecular battlefield, plant metabolic pathways are rapidly reprogrammed upon viral challenge to support defense responses. In this context, plants commonly suppress energy-consuming anabolic processes while enhancing catabolic pathways to meet increased energetic and redox demands (Llave, 2016). This metabolic reorganization is crucial for reshaping cellular micro-environment and thus transforms the cell into an optimal battlefield for plants to defend against viral replication.

Virus-induced metabolic reprogramming is associated with the alteration of different metabolites and leads to diverse impacts on host susceptibility. For instance, the levels of soluble sugars altered in the plant species infected with various virus strains (Love et al., 2005; López-Gresa et al., 2012; Fernández-Calvino et al., 2014). Changes in cellular carbohydrate status are associated with anti-viral responses. In tobacco plants, the upregulation of invertase activity leads to hexose accumulation, which is associated with resistance to viruses such as Tobacco mosaic virus (TMV) and Potato virus Y (PVY) (Herbers et al., 1996). Moreover, viral infection can modify sugar transport and allocation across the plant. Some viruses induce callose deposition at plasmodesmata and restrict sugar movement, which may eventually compromise virus spreading (De Storme and Geelen, 2014). Viral infection also significantly changes amino acid metabolism. The levels of amino acids increase in different plant species infected with Cucumber mosaic virus (CMV), Brome mosaic virus (BMV) and Tobacco rattle virus (TRV) (Xu et al., 2008). In Arabidopsis, knocking out DARK INDUCIBLE4 (DIN4), a key enzyme of branched-chain amino acid metabolism, significantly compromised TRV replication. This indicates that amino acid metabolism contributes to antiviral responses (Fernández-Calvino et al., 2014).

Cellular redox balance is important for the initiation of defense responses during viral infection. For example, mitochondrial metabolism plays a significant role in orchestrating metabolic reprogramming and associated defense responses during viral infections (Wang et al., 2022). It serves as a key mediator in the generation of reactive oxygen species (ROS) and the activation of phytohormone signaling pathways, which subsequently trigger downstream defense mechanisms, including programmed cell death, to impede viral proliferation (Colombatti et al., 2014). Furthermore, the interaction between α-ketoglutarate dehydrogenase and salicylic acid (SA) influences the mitochondrial electron transport chain, conferring basal defense against TMV (Liao et al., 2015). Chloroplast metabolism also plays a central role in plant defense mechanisms. Many viruses disrupt photosynthetic processes, leading to typical viral symptoms such as chlorosis and stunted growth (Zhao et al., 2016; Bhattacharyya and Chakraborty, 2018). This disruption impairs the chloroplast’s capacity to produce energy and generate ROS, both of which are vital for effective immune signaling (Caplan et al., 2015).

Several redox couples and thiol-based redox sensors are involved in maintaining cellular redox status, which contributes to plant defense responses (Pétriacq et al., 2016; Chae et al., 2023; Noctor et al., 2024). Glutathione, a tripeptide antioxidant, plays a central role in this process, with its reduced (GSH) and oxidized (GSSG) forms acting as a dynamic redox buffer. Elevated glutathione levels activate pathogenesis-related (PR) genes in tobacco and cucumber upon viral infection (Creissen et al., 1999; Gullner et al., 1999). The metabolic coenzyme, nicotinamide adenine dinucleotide (NAD), can enhance resistance to diverse pathogens by modulating the production of ROS and defense hormones, such as SA (Pétriacq et al., 2013). In Arabidopsis, elevated NAD levels are associated with increased resistance to *Pseudomonas syringae* and *Botrytis cinerea*, primarily through mitochondrial respiration-dependent ROS generation (Pétriacq et al., 2016), while the roles of NAD in anti-viral responses have not been investigated yet. Thiol-based redox sensors can transduce redox signals to regulate a variety of stress-adaptive mechanisms (Mou et al., 2003; Lindermayr et al., 2010). A well-documented example is the cytosolic thioredoxin-h5 (Trx-h5), which senses pathogen-induced redox changes and activates the master immune regulator, nonexpressor of pathogenesis-related genes 1 (NPR1) (Mou et al., 2003; Lindermayr et al., 2010). Upon pathogen attack, the accumulated ROS activates the expression of antioxidants and leads to a transiently reduced cellular environments. Under these conditions, the cytosolic Trx-h5 gains the reducing equivalents to resolve the disulfide bonds of NPR1 leading to the dissociation of NPR1 homotetramers. The resulting monomeric NPR1 then translocates into the nucleus to activate the expression of PR genes, thereby enhancing immunity against pathogens (Mou et al., 2003; Lindermayr et al., 2010).

Bamboo mosaic virus (BaMV) is a well-characterized positive-sense, single-stranded RNA virus of the *Potexvirus* genus. Its genome encodes five open reading frames (Lin et al., 1994) responsible for replication (Li et al., 1998; Li et al., 2001; Huang et al., 2004; Meng and Lee, 2017), movement (Chang et al., 1997; Lin et al., 2004; Liou et al., 2015) and encapsidation (Lan et al., 2010; Hung et al., 2014; DiMaio et al., 2015). To complete its infection cycle, BaMV exploits numerous host factors, including metabolic enzymes that play either proviral or antiviral roles. Notably, chloroplast phosphoglycerate kinase (PGK) and glyceraldehyde 3-phosphate dehydrogenase (GAPDH) have been shown to positively and negatively regulate BaMV replication, respectively (Lin et al., 2007; Prasanth et al., 2011; Cheng et al., 2013; Huang et al., 2017). Successful BaMV accumulation is tightly linked to chloroplasts and mitochondria— organelles that serve as central hubs for plant metabolism. By hijacking cytosolic enolase and mitochondrial voltage-dependent anion channel (VDAC), BaMV assembles a metabolon-like replication complex that bridges chloroplasts and mitochondria, thereby exploiting host energy and metabolic resources for its replication (Lin et al., 2007; Lee et al., 2022; Lin et al., 2025).

Although numerous studies have revealed the intricate association between plant metabolic processes and BaMV accumulation, most have focused on how BaMV exploits host factors to facilitate its replication. The mechanisms by which virus-induced metabolic reprogramming contributes to plant defense responses against viral infection remain insufficiently investigated. In this study, we investigated the regulatory roles of BaMV-induced metabolic reprogramming in defense responses in *N. benthamiana* plants. By integrating metabolic profiling, fluxomics, and proteomics, we demonstrated that BaMV infection reprograms plant metabolism by re-directing metabolic flux toward glycolysis, mitochondrial metabolism and amino acid biosynthesis. Notably, we revealed that mitochondrial NAD^+^-dependent malic enzyme 1 (NAD-ME1) modulates cytoplasmic redox homeostasis and defense signaling. Our findings highlight the central role of mitochondrial metabolism in orchestrating the defense response, shedding light on how the balance between host metabolic fluxes and redox states can influence the outcome of viral infections.

## Results

### Plants accumulate high levels of hexose phosphates and Krebs cycle intermediates in response to BaMV infection

To assess the impact of BaMV infection on host metabolic processes, we used liquid chromatography coupled with tandem mass spectrometry (LC-MS/MS) to quantify primary metabolites in BaMV-infected leaves. The results revealed significant increases in the levels of Glc-6-phosphate (G6P), Fru-6-phosphate (F6P) and Glc-1-phosphate (G1P) at 3 days post-infection (dpi). Following BaMV infection, the abundances of several Krebs cycle intermediates, including citrate, α-ketoglutarate, and succinate, were significantly increased, while the levels of fumarate and malate were dramatically decreased (Fig. 1; Supplementary Table S1). Given the close link between glycolytic and Krebs cycle intermediates and amino acid biosynthesis, we examined the levels of individual amino acids in BaMV-infected leaves. The infection induced dramatic changes in amino acid abundance, with notable increases in glutamate (Glu) and arginine (Arg), while valine (Val), aspartate (Asp), and threonine (Thr) levels were markedly reduced (Fig. 1; Supplementary Table S1). Other amino acids showed only minor changes in response to BaMV infection.

**Figure 1.**
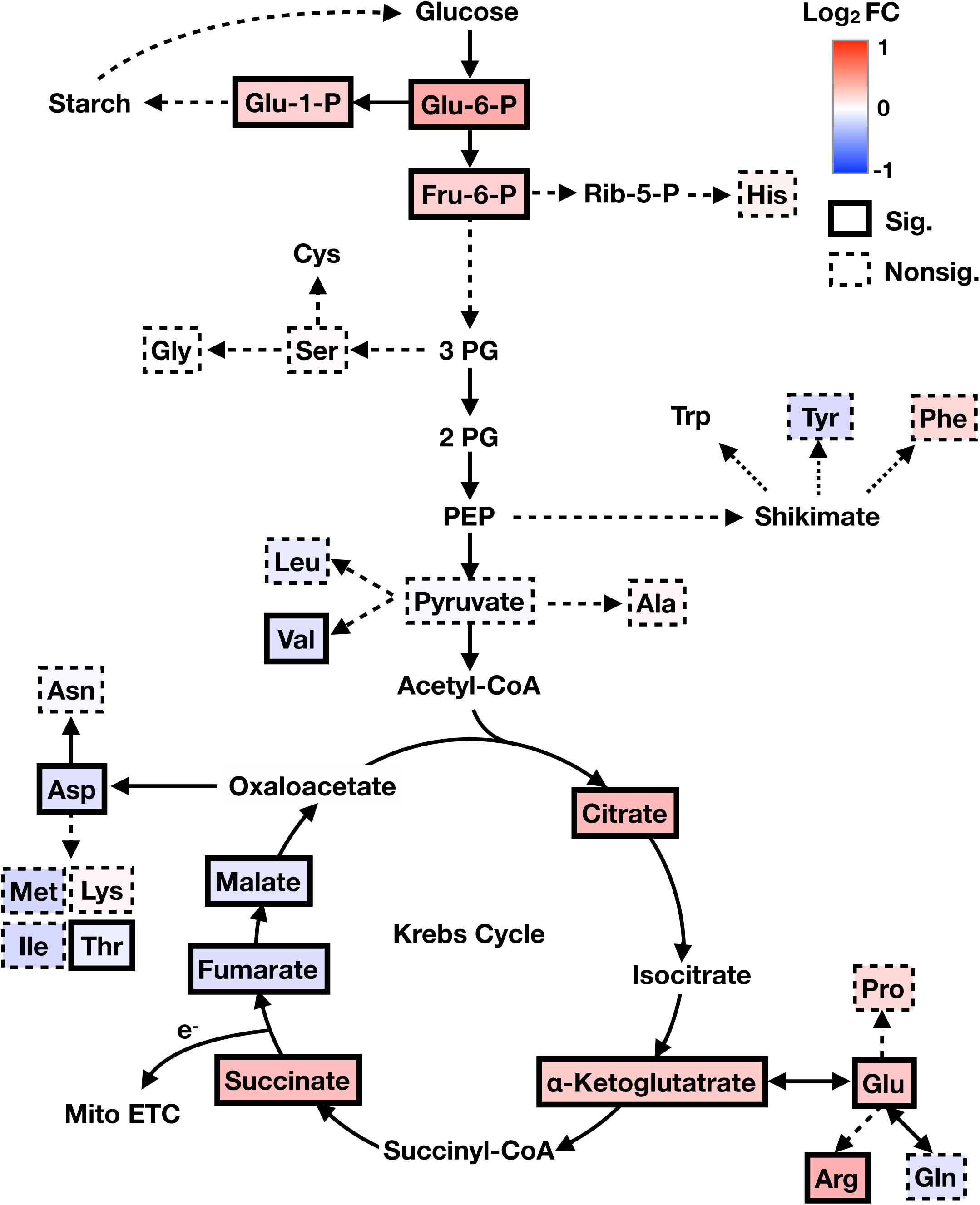
The changes of glycolytic, Krebs cycle intermediates and amino acid in BaMV-infected leaves. Leaves of *N. benthanmiana* plants were inoculated with 1 μg BaMV virions and harvested at 3 day-post-inoculation. Leaf samples were used for assaying metabolite levels using LC-MS/MS. The heatmaps summarize the log_2_ fold changes of metabolite levels in comparison to mock-infected leaves. Results are from 3 independent batches and each contained 4 to 5 biological replicates. Statistical analysis was conducted using unpaired t test with Holm-Šídák method (*P < 0.05). The symbol “Sig.” indicated significant changes, while “Nonsig.” Indicated non-significant changes.

To determine whether the changes in glycolytic and Krebs cycle metabolites were associated with carbohydrate metabolism, we further analyzed the accumulation of starch and major soluble sugars. In BaMV-infected leaves, the starch level was slightly increased, while the levels of Glc, Suc and maltose were significantly increased at 3 dpi (Supplementary Table S2). Since starch degradation usually operates with a low efficiency during the day, its contribution on the levels of soluble sugars is neglectable. Instead, the increase of soluble sugars in BaMV-infected leaves was likely attributed to the enhanced sugar transport from uninfected leaves. Thus, the increase in carbon pool in BaMV-infected leaves can further facilitate glycolysis and Krebs cycle.

### BaMV infection redirects metabolic flux toward glycolysis and the Krebs cycle thereby promoting amino acid biosynthesis

To monitor the dynamics of glycolysis and mitochondrial metabolism, we performed a metabolic flux analysis in BaMV-infected leaves. At 3 dpi, we fed BaMV-infected leaves with ^13^C-labeled Glc and analyzed ^13^C incorporation into metabolites using Liquid chromatography/quadrupole time-of-flight mass spectrometry (LC/Q-TOFMS). The ^13^C incorporation into glycolytic metabolites (G6P, F6P, G1P and pyruvate), as well as into Krebs cycle intermediates, such as succinate and malate, were significantly increased in BaMV-infected leaves compared to mock-treated leaves. Interestingly, the ^13^C incorporation into many amino acids including Ser, Gly, Phe, Ala, Val, Glu, Gln and Thr were also dramatically elevated upon BaMV infection (Fig. 2). These results indicate that BaMV infection reprograms metabolic flux to enhance glycolysis, the Krebs cycle, and associated amino acid biosynthesis.

**Figure 2.**
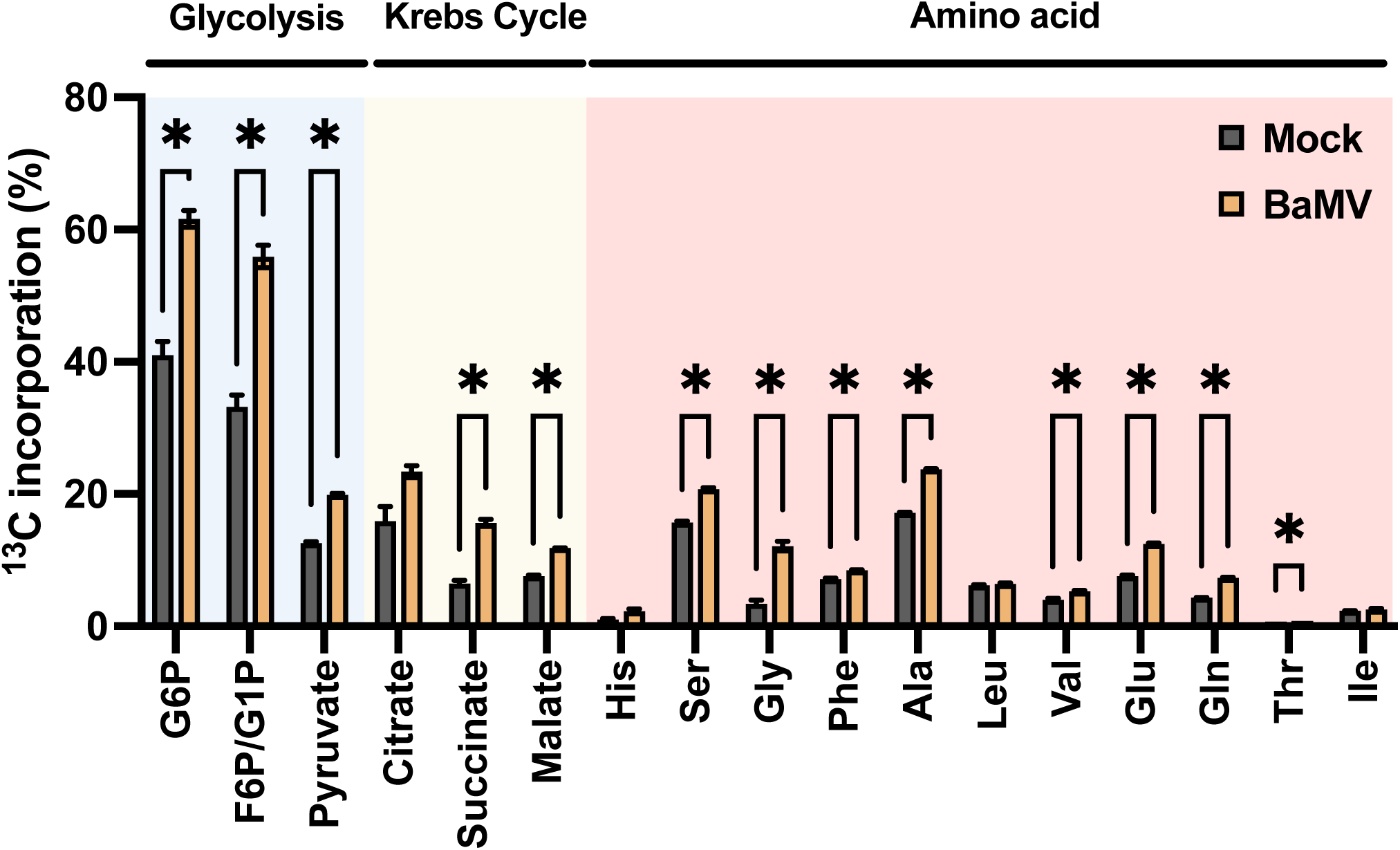
^13^C incorporation ratio of glycolytic, Krebs cycle intermediates and amino acid in leaves of mock- and BaMV-infected *N. benthamiana* plants. Leaves of *N. benthanmiana* plants were inoculated with 1 μg BaMV virions. At 3 day-post-inoculation, the infected leaves were fed with 15 mM of ^13^C-labeled glucose for 60 min. The leaves were harvested for metabolite extraction, and the ^13^C incorporation of respective metabolites were analyzed using LC/Q-TOFMS. The intensity of ^13^C-labeled metabolites was divided by the total intensity (labeled + unlabeled) to yield the ^13^C incorporation percentage. Results are the mean with SE from five biological replicates. Statistical analysis was conducted using Unpaired t test with Holm-Šídák method (*P < 0.05)

### BaMV infection dramatically downregulates photosynthetic proteins while upregulates mitochondrial enzymes

To identify host factors involved in promoting glycolysis and mitochondrial metabolism during BaMV infection, we conducted a time-resolved proteomic analysis in BaMV-infected leaves. We harvested mock- and BaMV-infected leaves for protein extraction and LC-MS/MS analysis at 1, 2, 3 dpi. Using this approach, we identified a total of 2,941 proteins. We considered proteins detected in at least three biological replicates and containing at least two unique peptides as valid targets. Applying this criterion, we selected 1,225 protein targets for further analysis. To obtain well-annotated protein information, we performed a homology search against the Arabidopsis proteome database. The principal component analysis (PCA) plot shows that mock samples (mock_1dpi, mock_2dpi and mock_3dpi) clustered close to each other, while BaMV-infected samples (BaMV_1dpi, BaMV_2dpi and BaMV_3dpi) clearly deviated from mock samples (Supplementary Fig. S1A). Notably, BaMV_3dpi samples formed a distinct cluster (Supplementary Fig. S1A), indicating that BaMV infection induces progressive and substantial changes in the host proteome over time.

To further evaluate the impact of BaMV infection on protein levels, we calculated the fold changes in abundance for each identified protein between BaMV-infected and mock samples. Based on these criteria, we identified 328 proteins that showed statistically significant changes in abundance (p < 0.05) during BaMV infection (Supplementary Table S3). The Venn diagram analysis indicated that the non-overlapping protein sets at 1, 2, and 3 dpi sample sets accounted for 25%, 24%, and 37% of the 328 differentially expressed proteins, respectively. Notably, fewer than 5% of the proteins overlapped across all three time points, and only four proteins showed significant changes at all stages of infection (Supplementary Fig. S1B). These results suggest that plant proteomes exhibit distinct dynamic changes throughout the course of BaMV infection.

Given that the 3-dpi sample set exhibited the most pronounced proteomic changes, we further analyzed those 156 differentially expressed proteins identified from this time point according to their biological functions (Supplementary Table S3). We identified 38 proteins associated with photosynthetic activity, 4 with cellular respiration, 9 with amino acid metabolism, 15 with other metabolic processes, 6 with stress response, 7 with redox homeostasis, and 17 with various other cellular functions. Furthermore, 45 proteins were classified under DNA, RNA and protein-related processes, while 15 remained uncharacterized with unknown function. Within photosynthesis and redox homeostasis categories, most identified proteins were less abundant upon BaMV infection (Fig. 3A and Supplementary Table S3), suggesting that virus infection impairs photosynthetic activity and disrupts cellular redox balance. In addition, the differential accumulation of enzymes, including Glu dehydrogenase (GDH1), Glu decarboxylase (GAD4), Gln synthetase (GSR1 and GSR2), and argininosuccinate synthase (ASS), may account for the accumulation of Glu and Arg in BaMV-infected leaves (Fig. 1 and Supplementary Table S3). Considering that BaMV infection leads to the upregulation in mitochondrial metabolism, we next examined the changes in mitochondrial proteins. Three key mitochondrial components involved in cellular respiration —cytochrome C oxidase 6b (COX6b-1), mitochondrial lipoamide dehydrogenase 1 (mtLPD1) and NAD-malic enzyme 1 (NAD-ME1) — were significantly increased in BaMV-infected leaves (Fig. 3B and Supplementary Table S3). These findings suggest that BaMV infection promotes the accumulation of mitochondrial key enzymes that boost respiratory activity.

**Figure 3.**
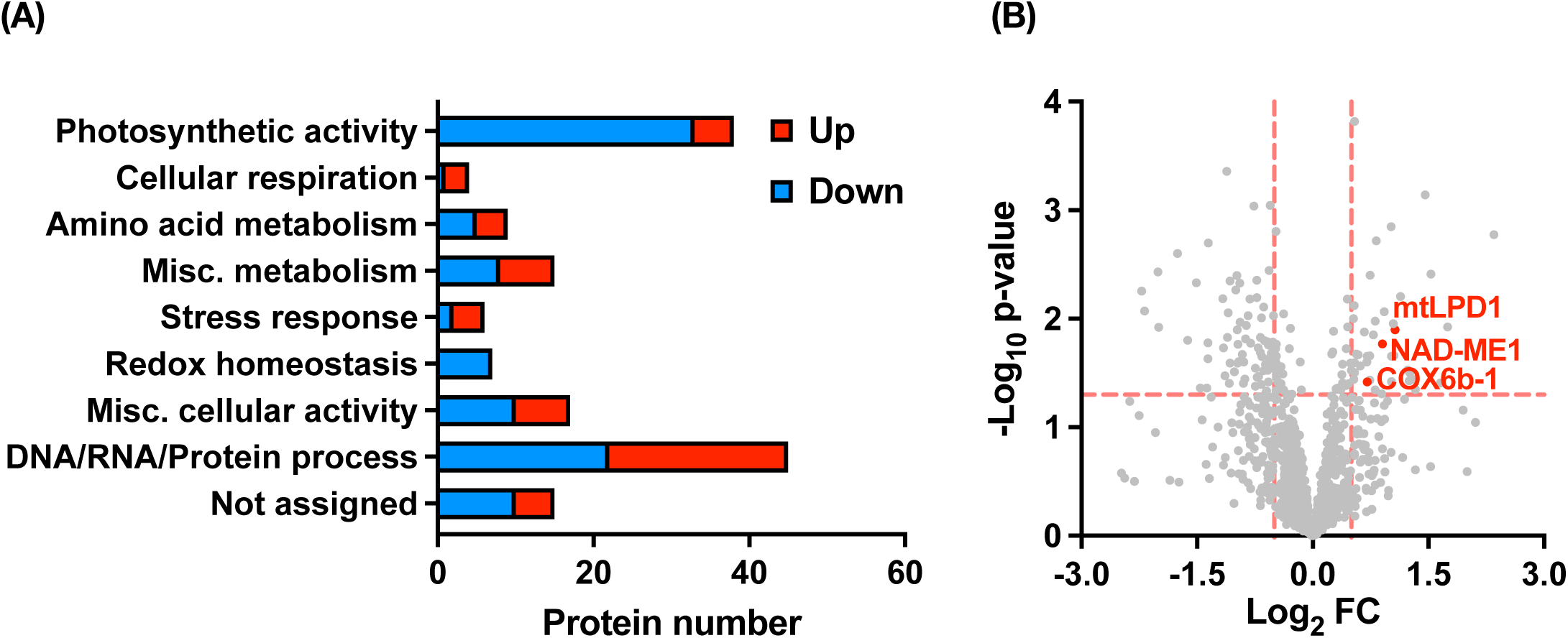
The proteome landscape in BaMV-infected leaves harvested at 3 day-post-inoculation. Leaves of *N. benthanmiana* plants were inoculated with 1 μg BaMV virions. At 3 day-post-inoculation, the infected leaves were harvested for protein extraction followed by a proteomic analysis. A) Functional categories of proteins showing significant changes in their abundance during BaMV infection. (B) Volcano plot indicating significantly increased mitochondrial targets upon BaMV infection. COX6b-1: cytochrome C oxidase 6b; mtLPD1: mitochondrial lipoamide dehydrogenase 1; NAD-ME1: NAD-malic enzyme 1. Results are from 3 to 4 biological replicates. Statistical analysis was conducted using unpaired t test (*P < 0.05).

### Silencing mtLPD1 and NAD-ME1 increases BaMV accumulation

To investigate the functional roles of identified mitochondrial components in the response to BaMV infection, we adopted an RNA interference approach to downregulate the expression of COX6b-1, mtLPD1 and NAD-ME1 in *N. benthamiana* leaves (Miki and Shimamoto, 2004). In the silenced leaves (hpCOX6b-1, hpmtLPD1 and hpNAD-ME1), the transcript levels of COX6b-1, mtLPD1 and NAD-ME1 were reduced to 12%, 28% and 16% of those in control leaves (EV; Fig. 4, A to C), respectively, indicating effective gene silencing. When examining BaMV accumulation, silencing COX6b-1 did not affect BaMV accumulation (Fig. 4A). However, silencing mtLPD1 and NAD-ME1 significantly increased BaMV titers to 1.5- and 2-fold of the levels in control leaves, respectively (Fig. 4, B and C). These results indicate that mtLPD1 and NAD-ME1 serve as pro-host factors that contribute to the plant’s defense against BaMV infection.

**Figure 4.**
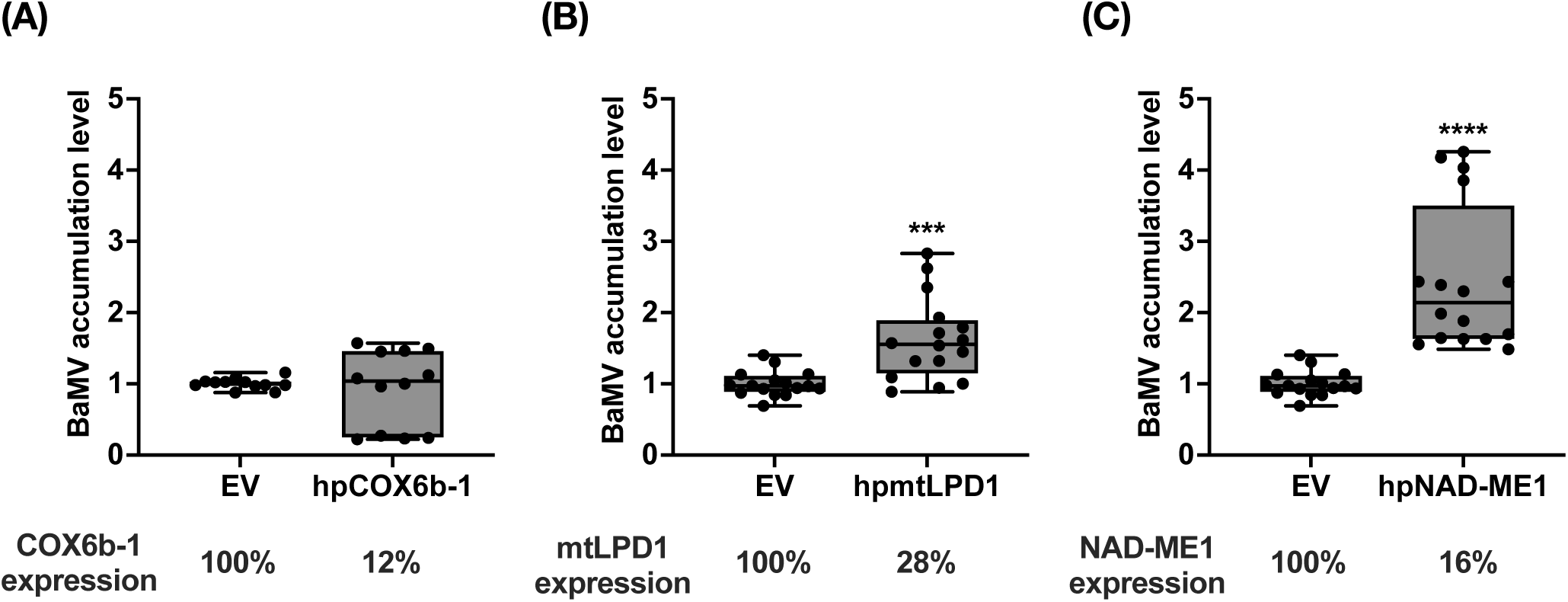
The impact of silencing mitochondrial targets on BaMV accumulation. Leaves of *N. benthanmiana* plants were infiltrated with agrobacteria harboring hairpin RNA silencing clones. At 3-day-post-infiltration, leaves were inoculated with 1 μg BaMV virions and harvested at 27-hour-post-inoculation. BaMV accumulation and target gene expression in silencing plants of A) COX6b-1, B) mtLPD1 and C) NAD-ME1 were detected using RT-qPCR. Results are from 12 to 16 biological replicates. Statistical analysis was conducted using unpaired t test (*P < 0.05; **P < 0.01; ***P < 0.001; ****P < 0.0001).

### Silencing NAD-ME1 increases the cytoplasmic NADH-to-NAD^+^ ratio and alters the expression patterns of defense-related genes

In Arabidopsis, NAD-ME1 is predicted to be a mitochondrial enzyme responsible for decarboxylating malate to generate pyruvate and NADH (Tronconi et al., 2008). To investigate whether the homolog of NAD-ME1 in *N. benthamiana* is also present in mitochondria, we fused *N. benthamiana* NAD-ME1 to an enhanced green fluorescence protein (eGFP) and performed agrobacteria-mediated transient expression in *N. benthamiana.* The confocal microscopic images showed that the signal of NAD-ME1-eGFP colocalized with the mitochondrial marker mito-mCherry (Supplementary Fig. S2), indicating that NAD-ME1 is a mitochondrial protein.

Mitochondrial metabolism is tightly associated with the balance of NAD(H) redox couple in the cytoplasm (Lim et al., 2020). To assess the impact of NAD-ME1 silencing on cytoplasmic NADH homeostasis, we employed a well-established NADH biosensor (Steinbeck et al., 2020). In both mock-treated and BaMV-infected leaves, silencing NAD-ME1 (hpNAD-ME1) led to a significant increase in the cytoplasmic NADH-to-NAD^+^ ratio compared to control leaves at 3 dpi (EV; Fig. 5, A and B). This indicates disrupting NAD-ME1 expression may affect mitochondrial metabolism and perturbs the cytoplasmic NAD(H) redox balance. Interestingly, BaMV infection alone slightly elevated the NADH-to-NAD^+^ ratio compared to mock-treated leaves (Fig. 5B), suggesting that BaMV-induced accumulation of NAD-ME1 only partially affects cytoplasmic NADH homeostasis (Fig. 3B). This implies that additional host factors likely act in concert with NAD-ME1 to maintain NAD(H) redox balance during BaMV infection.

**Figure 5.**
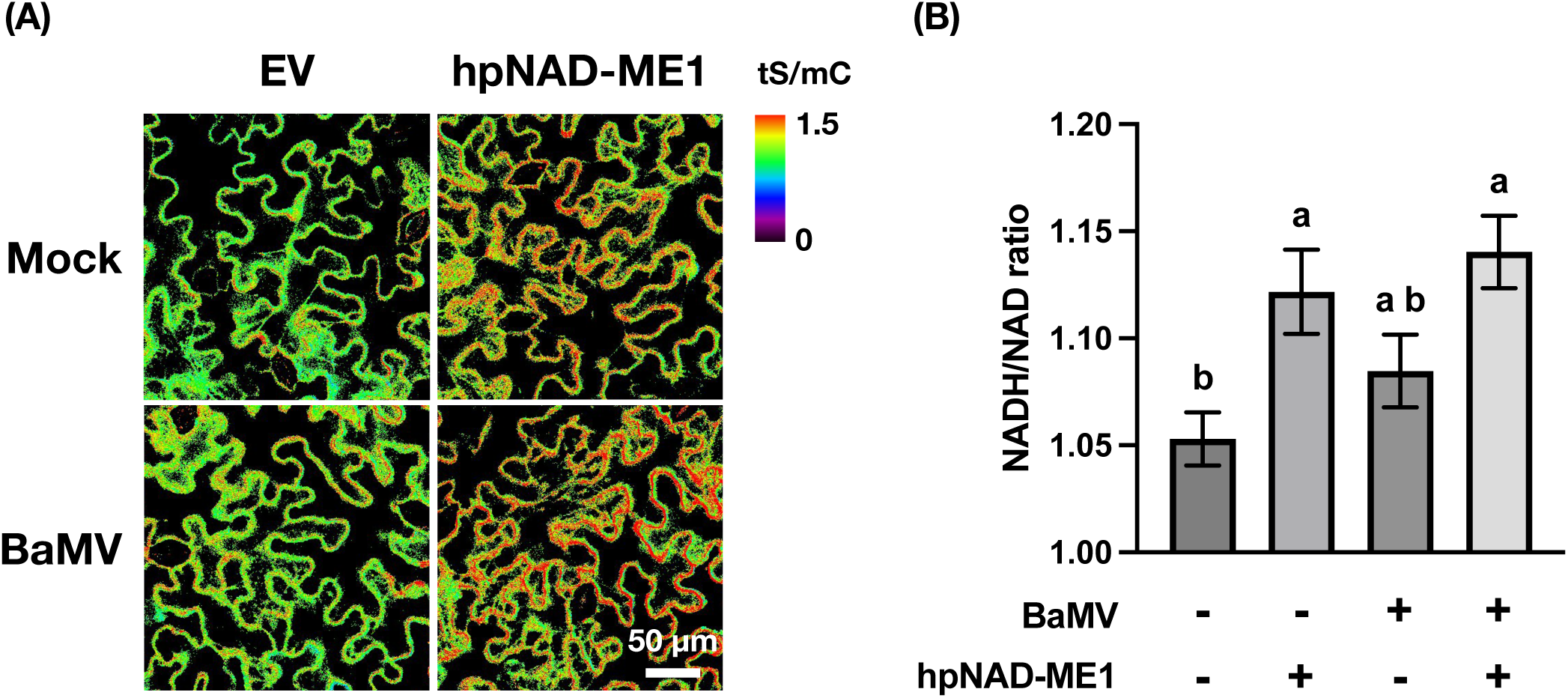
The impact of silencing NAD-malic enzymes 1 (NAD-ME1) on cytoplasmic NADH-to-NAD ratio. Leaves of *N. benthanmiana* plants were infiltrated with agrobacteria harboring NAD-ME1 silencing clones, NADH biosensor and BaMV infectious clone. At 3-day-post-infiltration, leaves were subjected to confocal microscopy. The florescence of tSapphire (tS, NADH binding module) and mCherry (mC, control module) were recorded sequentially. The intensity of tS was divided by the intensity of mC to yield the ratio representing NADH-to-NAD ratio. A) The confocal images representing the cellular NADH-to-NAD ratio. B) The qualitative results of the cellular NADH-to-NAD ratio. Results are from 18 biological replicates. Statistical analysis was conducted using one-way ANOVA with Tukey’s post-hoc test (P < 0.05).

Cellular redox balance can affect several phytohormone-mediated defense responses during pathogen infection (Pétriacq et al., 2016; Chae et al., 2023; Noctor et al., 2024). To investigate whether the redox imbalance resulting from the disruption of NAD-ME1 expression can affect defense responses during viral infection, we examined the expression profile of defense marker genes (Wang et al., 2000; Lorenzo et al., 2003; Lee et al., 2013a). In BaMV-infected leaves, the levels of PR2b, NPR1 and RBOHB were significantly increased (Fig. 6, A, B and F), while the levels of LOX, PDF1.2, ERF1 remained comparable to mock-treated leaves (Fig. 6, C, D and E). Silencing NAD-ME1 dramatically changed the expression pattern of defense genes. In mock-treated leaves, silencing NAD-ME1 up-regulated the expression of PR2b (Fig. 6A) while down-regulated the expression of NPR1, LOX and ERF1 (Fig. 6, B, C and E). In BaMV-infected leaves, silencing NAD-ME1 further elevated the levels of PR2b, PDF1.2 and RBOHB (Fig. 6, A, D and F). Notably, the expression of NPR1 remained lower in NAD-ME1 silencing leaves compared to control leaves during viral infection (Fig. 6B). Collectively, silencing NAD-ME1 alters the expression profile of defense genes, likely due to an imbalance in the cytoplasmic NAD(H) redox couple. Since NPR1 activation is tightly associated with cytoplasmic redox dynamics (Mou et al., 2003; Lindermayr et al., 2010), we further examined the impact of NAD-ME1 silencing-induced redox imbalance on NPR1 activation. Interestingly, in the NAD-ME1-silenced leaves, the levels of exogenously expressed NPR1-GFP oligomers and monomers were dramatically reduced compared to the control leaves, as observed under both reducing and non-reducing conditions (Supplementary Fig. S3A). Furthermore, silencing NPR1 significantly increased BaMV accumulation (Supplementary Fig. S3B). These results indicate that NAD-ME1 silencing-induced redox imbalance may compromise NPR1 protein stability and interfere with its function in regulating downstream defense responses during BaMV infection.

**Figure 6.**
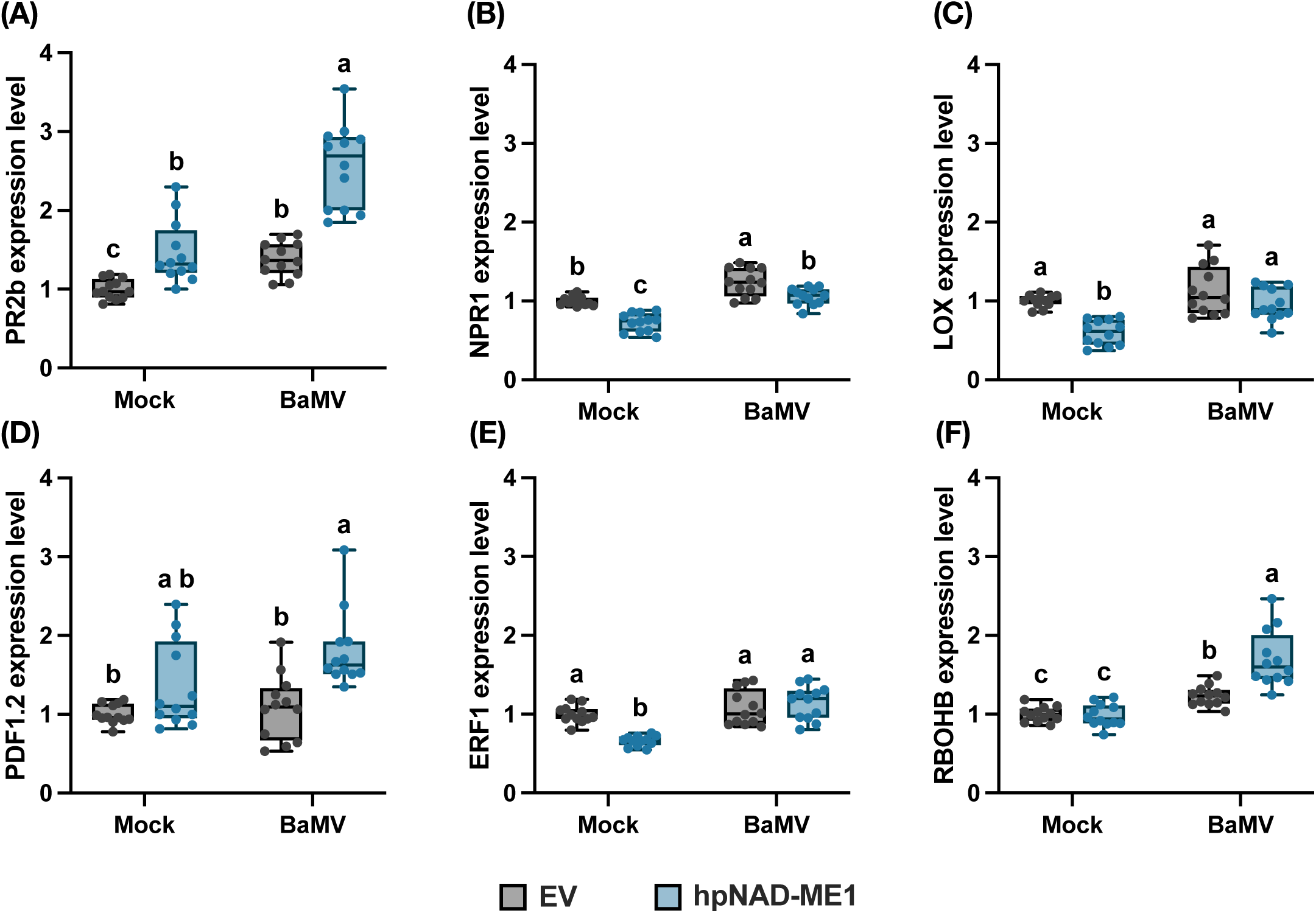
The expression profile of defense genes in the silencing plants of NAD-malic enzymes 1. Leaves of *N. benthanmiana* plants were infiltrated with agrobacteria harboring hairpin RNA silencing clones. At 3-day-post-infiltration, leaves were inoculated with 1 μg BaMV virions and harvested at 27-hour-post-inoculation. Defense gene expression of A) PR2b, B) NPR1, C) LOX, D) PDF1.2, E) ERF1 and E) RBOHB were detected using RT-qPCR. Results are from 12 to 16 biological replicates. Statistical analysis was conducted using one-way ANOVA with Tukey’s post-hoc test (P < 0.05).

## Discussion

Viral infection often induces substantial alterations in host metabolism, redirecting cellular resources and modifying the intracellular environment to support viral replication. At the same time, these metabolic changes can also activate host defense responses against the invading virus. However, the molecular mechanism underlying metabolic reprogramming contributing to defense responses remains poorly understood. In this study, we incorporated multifaceted omic approaches and genetic validation to systemically investigate how plants regulate their metabolic machinery to fine-tune defense responses during viral infection. By analyzing the metabolome and fluxome in *N. benthamiana* plants infected with BaMV, we demonstrated that plants rewired their metabolic flux toward glycolysis, mitochondrial metabolism and the biosynthesis of amino acids (Fig. 1; Fig. 2). Proteomic analysis of BaMV-infected leaves revealed a significant decrease in photosynthetic proteins and an increase in mitochondrial metabolic enzymes, including COX6b-1, mtLPD1 and NAD-ME1. This indicates that mitochondrial proteins play key roles in coordinating metabolic reprogramming during viral infection (Fig. 3). By analyzing the NAD-ME1-silenced leaves in response to BaMV infection, we found that silencing of NAD-ME1 increased cytoplasmic NADH-to-NAD^+^ ratio, altered the expression profile of defense-related genes, and led to higher BaMV accumulation (Fig. 4C; Fig. 5; Fig.6). Together, these results highlight the pivotal role of NAD-ME1 in fine-tuning antiviral defense responses through maintaining cellular redox balance during infection.

### Plants undergo a source-to-sink transition upon viral infection

Our fluxomic study showed that BaMV infection prompts a pronounced reprogramming of host central metabolism, characterized by a redirection of metabolic flux toward glycolysis, mitochondrial activity, and amino acid biosynthesis (Fig. 2). This metabolic shift was accompanied by an accumulation of soluble sugars, glycolytic intermediates, and Krebs cycle metabolites (Fig. 1; Supplementary Table S2), reflecting increased metabolic demand in infected tissues. These findings are consistent with previous studies showing elevated sugar levels in Tomato mosaic virus-infected tomato and TRV-infected Arabidopsis plants (López-Gresa et al., 2012; Fernández-Calvino et al., 2014). Such metabolic responses are believed to enhance sink strength in infected tissues, thereby promoting the mobilization and import of carbohydrates from photosynthetically active source leaves to support the high energy demands of biosynthesis, defense processes and viral replication (Llave, 2016). This carbon enrichment dramatically alters the balance of cellular nitrogen-to-carbon ratio, potentially triggering transcriptional reprogramming of metabolic pathways through feedback regulation (Zheng, 2009). Supporting this notion, our proteomic data revealed a widespread reduction of photosynthesis-associated proteins in BaMV-infected leaves (Fig. 3; Supplementary Table S3). Beyond a passive consequence of stress, this repression may also reflect active viral interference with chloroplast function, as BaMV has been shown to hijack chloroplast-localized host factors such as PGK and photosystem II oxygen-evolving complex proteins to facilitate its replication (Lin et al., 2007; Cheng et al., 2013; Huang et al., 2021). Suppression of photosynthetic activity likely amplifies the source-to-sink transition by reinforcing carbohydrate import into infected tissues. Interestingly, experimental evidence suggests that increased sink strength is linked to enhanced antiviral defense. For example, *Nicotiana* lines overexpressing cell-wall invertase, which drives increased sugar accumulation, display enhanced resistance to viral pathogens (Herbers et al., 1996). Similarly, virus-resistant tomato cultivars infected with Tomato yellow leaf curl virus accumulate higher levels of amino acids and related metabolites compared to susceptible lines (Sade et al., 2015). Taken together, these findings suggest that source-to-sink transition is not merely a passive consequence of infection but may instead represent an active antiviral defense mechanism by reallocating metabolic resources and modulating defense signaling in infected tissues.

### Glu and Arg act as potential signaling molecules in antiviral defense

Although our fluxomic analysis revealed a general upregulation of amino acid biosynthesis in BaMV-infected leaves, the accumulation patterns of individual amino acids were notably diverse (Figs. 1 and 2). This metabolic heterogeneity suggests that plants may selectively modulate specific amino acid pools in response to viral infection, possibly reflecting distinct metabolic demands or defense priorities. Among the amino acids with significant changes, the elevated levels of Glu and Arg, along with the differential expression of associated metabolic enzymes (GDH1, GAD4, GSR1, GSR2, and ASS), are particularly noteworthy (Fig. 1 and Supplementary Table S3). Glu and Arg are increasingly recognized for their roles in plant defense against biotic stresses (Seifi et al., 2013; Winter et al., 2015; Qiu et al., 2020). Exogenous application of Glu can induce defense gene expression and enhance resistance against fungal pathogens both in pear and tomato plants (Jin et al., 2019; Sun et al., 2019). Similarly, Arg pretreatment reduces *Botrytis cinerea* lesion development in tomato fruits, likely by activating defense-related enzymes (Zheng et al., 2011). In addition, Arg is also the metabolic precursor for polyamines and nitric oxide, two important signaling molecules involved in a wide range of stress responses (Winter et al., 2015). Given these functional roles, it is likely that the upregulation of Glu and Arg metabolism contributes to antiviral defense responses, even though their accumulation patterns may vary across different plant-virus interactions (Blua et al., 1994; Fernández-Calvino et al., 2014). The enhanced levels of these amino acids in BaMV-infected leaves may thus reflect an adaptive reprogramming of nitrogen metabolism to reinforce host immunity.

### NAD-ME1 coordinates mitochondrial metabolism and maintains the balance of cytoplasmic NAD(H) redox couples during BaMV infection

Our proteomic study shows that the abundance of mitochondrial protein NAD-ME1 was significantly increased in BaMV-infected leaves (Fig. 3B). Subsequent loss-of-function experiments demonstrated that silencing mitochondrial NAD-ME1 led to an increased cytoplasmic NADH-to-NAD^+^ ratio (Fig. 5), suggesting its role in maintaining cytoplasmic redox balance under viral stress. In mitochondria, NAD-ME1 catalyzes the decarboxylation of malate to pyruvate, thus facilitating carbon flux into the Krebs cycle and supporting mitochondrial respiration (Grover and Wedding, 1982). Consistently, knocking out of mitochondrial NAD-ME in *Arabidopsis* causes oscillatory fluctuations in Krebs cycle intermediates (Tronconi et al., 2008). Furthermore, pharmacological inhibition of mitochondrial NADH consumption has been shown to elevate cytoplasmic NADH and NADPH levels, further supporting the direct influence of mitochondrial metabolism on cytosolic redox status (Lim et al., 2020). In this context, silencing NAD-ME1 likely compromises mitochondrial metabolism, leading to cytoplasmic NADH accumulation and disruption of redox homeostasis under viral stress. Intriguingly, BaMV appears to exploit host metabolism by hijacking cytosolic enolase and mitochondrial VDAC to assemble a metabolon-like replication complex (Lee et al., 2022; Lin et al., 2025). This structure not only facilitates efficient viral replication but may also promote mitochondrial clustering and metabolic reprogramming, further perturbing cytoplasmic redox balance, and potentially weakening the host’s defense responses.

### The dynamics of cytoplasmic NAD(H) redox couples play a crucial role in defense signaling during BaMV infection

In addition to prompting cytoplasmic reduction, silencing mitochondrial NAD-ME1 also significantly increased BaMV accumulation and altered the transcriptional landscape of defense genes (Figs. 4C and 6). These results suggest that persistent cytoplasmic reduction may interfere with redox-sensitive defense signaling during BaMV infection. One key example is the SA-dependent immune regulator NPR1, whose activation is tightly governed by the cellular redox state (Mou et al., 2003; Lindermayr et al., 2010). Under normal conditions, NPR1 exists as inactive oligomers in the cytosol, stabilized by disulfide bonds. Upon pathogen challenge, a transient increase in ROS triggers antioxidant responses and facilitates localized reduction. This redox shift enables cytosolic Trx-h5 to cleave disulfide bonds in NPR1, allowing its monomeric forms to translocate into the nucleus and activate downstream defense genes (Mou et al., 2003; Lindermayr et al., 2010). However, excessive and prolonged cytoplasmic reduction, such as that induced by NAD-ME1 silencing, may disrupt this finely tuned redox cycling, thereby hindering NPR1 activation and transcriptional defense reprogramming. This redox imbalance could partially explain the dysregulation of defense gene expression and the corresponding increase in BaMV titers observed in NAD-ME1-silencing leaves (Fig. 4C). It is well-documented that reduced protein forms are more susceptible to irreversible oxidative damage and proteolytic degradation under stress conditions (Pajares et al., 2015). Supporting this notion, we observed a marked reduction in the accumulation of exogenously expressed NPR1-GFP in NAD-ME1-silenced leaves (Supplementary Fig. S3A), further implicating the redox imbalance in compromising the stability of defense-related proteins. Interestingly, a reductive cellular environment has previously been shown to favor BaMV replication (Chen et al., 2013), indicating that cellular redox status can directly influence viral replication. Collectively, these findings highlight the critical role of redox homeostasis in coordinating immune signaling, stabilizing key defense components, and restricting viral replication.

## Materials and methods

### Plant material and growth conditions

The *N. benthamiana* plants were grown in a climate chamber equipped with regular white light tubes. The light intensity was set as 120 μmol photons m^-2^ s^-1^ with an 8-h dark/16-h light regime, and the temperature was set as 28°C. The forth leaves of 21-day-old plants were used for further analyses.

### Construct generation

To monitor the subcellular localization of NAD-ME1, the construct, pBIN61-MAD-ME1-eGFP-HA, was generated. The full-length coding region of NAD-ME1 was amplified from the cDNA library of *N. benthamiana* plants. The NAD-ME1 amplicons were subject to restriction enzyme treatment using XhoI, and ligated to the vector, pBIN61-eGFP-HA, linearized by XhoI and PmlI. To generate the clones, pBIN61-mito-mCherry, for visualization of mitochondria, the signal peptide of an ATPase gene (Lee et al., 2013b) was amplified from the cDNA library of *Arabidopsis thaliana* plants. The amplicons were by XbaI and BamHI, and ligated into the vector, pBIN61-3HA-mCherry linerizaed by XbaI and BamHI. The construct, pBIN61-NbNPR1-eGFP-HA, was generated to test NPR1 protein stability. The full-length coding region of NbNPR1 was amplified from the cDNA library of *N. benthamiana* plants. The NbNPR1 amplicons were digested by XhoI, and ligated to the vector, pBIN61-eGFP-HA, linearized by XhoI and PmlI. To perform the gene silencing assays, the clones, pANDA-NAD-ME1, pANDA-mtLPD1, pANDA-COX6b-1 and pANDA-NbNPR1, were generated. Approximate 200-to-300 bp fragments of the above genes were amplified from the cDNA library of *N. benthamiana* plants. The DNA fragments were incorporated into the vector pCR™8/GW/TOPO™ (Invitrogen^TM^) to generate the subclones, pCR8-NAD-ME1, pCR8-mtLPD1, pCR8-COX6b-1 and pCR8-NbNPR1. The DNA segments of these subclones were further incorporated into the destination vector, pANDA (Miki and Shimamoto, 2004), using Gateway™ LR Clonase™ (Invitrogen^TM^). All primer pairs were listed in Supplementary Table S4.

### RNA extraction and RT-qPCR

The *N. benthamiana* leaf was frozen in liquid nitrogen and ground into fine powder using mortar and pestle. Approximate 100 mg of leaf powder was used for RNA extraction by utilizing TRIzol™ Reagent (Invitrogen^TM^). The RNA concentration was quantified by using NanoDrop spectrometer (ND-2000, Thermo Fisher Scientific). One microgram of total RNA was used for cDNA library synthesis by utilizing TOOLSQuant II Fast RT Kit (KRT-BA06, TOOLS). The synthesized cDNA samples were diluted 10 times using nuclease-free water, and 4.5 μL of diluted cDNA samples was mixed with 0.25 μL of 10 μM forward primer, 0.25 μL of 10 μM reverse primer and 5 μL of *Power* SYBR^TM^ Green PCR Master Mix (Applied Biosystems^TM^). The PCR reaction was conducted in the thermocycler (12k Flex, QuantStudio™ 3, Applied Biosystems^TM^). The calculation of gene expression was according to the method of 2^-(ΔΔCt)^ (Winer et al., 1999; Schmittgen et al., 2000). All qPCR primer pairs were listed in Supplementary Table S4.

### Starch and soluble sugar quantification

Twenty milligrams of pulverized leaf tissue was mixed with 500 μL of 80% (v/v) ethanol and incubated at 90°C for 30 min. The mixture was centrifuged at 20,000 X *g* for 15 min at ambient temperature. The supernatant was transferred into a microtube, while the pellet was resuspended in 250 μL of 50% (v/v) ethanol and incubated at 90°C for 30 min. The mixture was centrifuged at 20,000 X *g* for 15 min at ambient temperature. The supernatant was transferred into the previous microtube and used for the quantification of soluble sugars, while the pellet was dehydrated and used for the quantification of starch. The detailed detection method was described in the previous publication (Hou et al., 2019).

### Metabolomic analysis

One hundred milligrams of pulverized leaf tissue was mixed with 500 μL of methanol containing 5 ppm of succinate-2,2,3,3-d4 as internal standard. The mixture was incubated at 4°C for 10 min followed by the addition of 200 μL of chloroform and 500 μL of double-distilled water. After vigorous vortex at 4°C for 5 min, the mixture was subject to centrifugation at 20,000 X *g* for 15 min at 4°C. The aqueous phase was transferred into a microtube followed by dehydration using a vacuum centrifugal concentrator (ScanSpeed MaxiVac, ScanVac). The pellet was resuspended in 100 μL of 50% methanol.

For the targeted metabolomic analysis, the extract was separated at 30oC using an Agilent InfinityLab Poroshell 120 HILIC column (2.7 μm, 2.1 × 100 mm) connected to a Thermo Scientific Vanquish Horizon UHPLC System and subjected to a Thermo Scientific Orbitrap Fusion Lumos Tribrid Mass Spectrometer. The mobile phases were composed of eluent A (50% [v/v] acetonitrile) and eluent B (90% [v/v] acetonitrile). Both eluents were supplemented with 20 mM ammonium acetate (pH 9.0). The flow rate was set as 500 μL/min. The mass spectrometer was equipped with an electrospray ionization (ESI) source operating in negative full-scan ion mode to collect signals ranging from 70 to 1000 (m/z). The chromatogram acquisition, detection of mass spectral peaks, and their waveform processing were performed using Xcalibur^TM^ software (Thermo Fisher Scientific).

For the metabolic flux analysis, the metabolite extract was separated at 40°C using an ACQUITY UPLC BEH amide column (1.7 μm, 2.1 × 100 mm, Waters Corp., Milford, MA, USA) connected to an Agilent 1290 II Infinity Ultra-High-Performance Liquid Chromatography system (Agilent Technologies, Palo Alto, CA, USA) and subjected to an Agilent 6545XT quadrupole time-of-flight (Q-TOF) mass spectrometer (Agilent Technologies, Palo Alto, CA, USA). The mobile phases were composed of eluent A (deionized water) and eluent B (90% [v/v] acetonitrile). Both eluents were supplemented with 15 mM ammonium acetate and 0.3% NH_3_·H_2_O. The flow rate was set as 300 uL/min. The mass spectrometer was equipped with an Agilent Dual AJS electrospray ionization (ESI) source operating in positive and negative full-scan ion mode to collect signals ranging from 60 to 1500 (m/z). The chromatogram acquisition, detection of mass spectral peaks, and their waveform processing were performed using Agilent Qualitative Analysis 10.0 (Agilent, USA). Isotopologue extraction and the correction of natural isotope abundance was performed using Agilent Profinder 10.0 (Agilent, USA). The mass tolerance and retention time tolerance were set to ±10 ppm and ±0.5 minutes, respectively.

### Protein extraction and proteomic analysis

Approximate 25 mg of leaf powder was mixed with 250 μL of extraction buffer (20 mM Tris-HCl, pH 7.5, 5 mM EDTA, 10 mM NaCl, 8 M Urea, 10 mM DTT) and incubated for 10 min at 37°C with mild shaking. The extract was subject to centrifugation at 20,000 X *g* for 10 min at 4°C to remove leaf debris. Two hundred microliter of supernatant was mixed with 4 volumes of absolute acetone and subjected to centrifugation at 20,000 X *g* for 10 min at 4°C to precipitate protein. The protein pellet was washed with 80% of acetone and dehydrated under a chemical hood. The dehydrated pellet was resuspended in 50 μL of extraction buffer. The protein concentration was determined using Pierce™ 660nm Protein Assay Reagent (Thermo Scientific™).

Protein digestion in the S-Trap microcolumn was performed according to the protocol of S-Trap-IMAC with some modifications (Chen et al., 2023; Chen et al., 2024). In brief, 10 µg of protein was reconstituted in the buffer containing 4M urea and 5% (v/v) sodium dodecyl sulfate. The protein sample was reduced and alkylated using 10 mM Tris(2-carboxyethyl)phosphine (TCEP) and 40 mM chloroacetamide (CAA) at 45°C for 15 min, respectively. A final concentration of 5.5% (v/v) PA was applied to the protein sample followed by mixing with a six-fold volume of binding buffer containing 90% (v/v) methanol in 100 mM of triethylammonium bicarbonate (TEAB). After moderate vortexing, the solution was loaded into an S-Trap microcolumn and subject to centrifugation at 4,000 X *g* for 1 min. The column was washed with 350 µL binding buffer three times. Finally, 20 µL of digestion solution (0.2-unit Lys-C and 200 ng trypsin in 50 Mm TEAB) was added to the column and incubated at 47°C for 2 h. Each digested peptide was eluted using 40 µL of three buffers consecutively: (1) 50 Mm TEAB, (2) 0.2% (v/v) formic acid in H_2_O, (3) 50% (v/v) acetonitrile. Eluted solutions were collected in a tube and dehydrated under vacuum. Dried peptides were suspended in 50 µL H_2_O. Five micrograms of peptides were desalted using Evotip and dehydrated under vacuum.

Peptide samples were loaded onto Evotips Pure and eluted into an EV1137 performance column (15 cm x 150 µm ID, 1.5 µm, Evosep Biosystems) on the Evosep One LC system (Evosep Biosystems) connected to a timsTOF HT mass spectrometer (Bruker Daltonics). The mass spectrometer was set to PASEF scan mode for data-dependent acquisition. All spectra were acquired over an m/z range of 100 to 1,700 with 10 PASEF ramps. The TIMS ranges were initially set from 0.6 to 1.6 1/K0 [V·s/cm²] with settings of 100 ms ramp and accumulation time (100% duty cycle) and a ramp rate of 9.43 Hz; this resulted in a total cycle time of 1.17 s. Singly charged precursors were excluded based on their position in the m/z-IM plane using a polygon shape, and linear precursor repetitions were set at a target intensity of 20,000 with a threshold of 2,500. Active exclusion was enabled with a 0.4 min release time. The collision energy remained at default, with a base of 1.6 1/K0 [V·s/cm²] set at 59 eV and 0.6 1/K0 [V·s/cm²] set at 20 eV. Isolation widths were set to 2 m/z at <700 m/z and 3 m/z at >800 m/z. The TIMS ranges were initially set from 0.6 to 1.6 1/K0 [V·s/cm²].

These raw files were searched with SpectroMine™ (version SpectroMine 4.2.230428.52329, Biognosys) against the established proteome database of N. benthamiana (Kourelis et al., 2019) using the following settings: fixed modification: carbamidomethyl (C); variable modifications: acetyl (protein N-term), oxidation (M); enzyme: trypsin and LysC with up to two missed cleavages. Mass tolerances were automatically determined by SpectroMine, and other settings were set to default parameters. Identified results were filtered by a 1% FDR at the precursor, peptide, and protein levels.

### SDS-PAGE and immunoblotting

Protein extraction followed the approach mentioned above. For the non-reducing condition, DTT was omitted in the extraction buffer. The protein samples were assayed by SDS-PAGE using 8% (v/v) acrylamide gel and immunoblotting. The signals of actin and NPR1-GFP were detected using commercial actin antibody (A0408, Sigma-Aldrich) and GFP-tag antibody (CPA9023, Cohesion Bioscience), respectively.

### Confocal laser scanning microscopy

The leaf disc of *N. benthamiana* plants was placed onto a glass slide together with a few droplets of double distilled water followed by covering with a coverslip. The fluorescence signals were monitored using a confocal microscope (Olympus FV3000). The excitation wavelengths were set at 488 nm for eGFP, 561 nm for mCherry and 405 nm for tSapphire (NADH biosensor). The emitted signals of eGFP, mCherry and tSapphire were collected at 510 ± 10 nm, 610 ± 10 mn and 515 ± 10 nm, respectively.

### Accession numbers

Sequence information in this article can be retrieved in the data library of Sol Genomic Network and QUT under the below accession numbers: NAD-ME1: Nbv5.1tr6319150; mtLPD1: Niben101Scf01334g06004.1; COX6b-1: Nbv5.1tr6218676; NPR1: Niben101Scf14780g01001.1. The curated sequences are listed in Supplementary Table S4.

## Acknowledgments

We are grateful to the specialists at the core laboratories of the Institute of Plant and Microbial Biology (IPMB), Academia Sinica, for their support in various experiments: Ms. Yu-Ching Wu of the Small Molecule Metabolomics Core Lab for assistance with metabolomic assays; Dr. Chuan-Shih Hsu of the Proteomics Core Lab for proteomic analyses; Ms. Mei-Jane Fang and Mr. Ji-Ying Huang of the Cell Biology Core Lab for assistance with cell biology experiments; and Dr. Wen-Dar Lin of the Bioinformatics Core Lab for bioinformatic analysis. Special thanks go to Dr. Chih-Yu Lin and Mr. Gong-Min Lin of the Metabolomics Core Laboratory at the Agricultural Biotechnology Research Center for their assistance with metabolic flux analyses. We also thank Ms. Li-Shih Liao for preparing *N. benthamiana* plants and Dr. Shu-Chuan Lee for assistance with clone construction in gene silencing experiments.

## Supplemental data

**Supplementary Fig. S1.** The landscape of proteome between mock- and BaMV-infected leaves.

**Supplementary Fig. S2.** The subcellular localization of NAD-malic enzymes 1 (NAD-ME1).

**Supplementary Fig. S3.** The impact of silencing NAD-malic enzymes 1 (NAD-ME1) on protein stability of nonexpressor of pathogenesis-related genes 1 (NPR1) and the impact of silencing NPR1 on BaMV accumulation.

**Supplementary Table S1.** The levels of glycolytic, Krebs cycle intermediates and amino acids in mock- and BaMV-infected leaves at 3 day-post-inoculation.

**Supplementary Table S2.** The levels of starch and soluble sugars in leaves of mock- and BaMV-infected *N. benthamiana* plants.

**Supplementary Table S3.** Proteins displaying significant changes in abundance in BaMV-infected leaves compared to mock-infected leaves at 3 day-post-inoculation.

**Supplementary Table S4.** Sequences of primers and genes in this study.

