## Supplemental Figures for "Virus-induced upregulation of mitochondrial metabolism modulates cytosolic redox balance and defense responses"

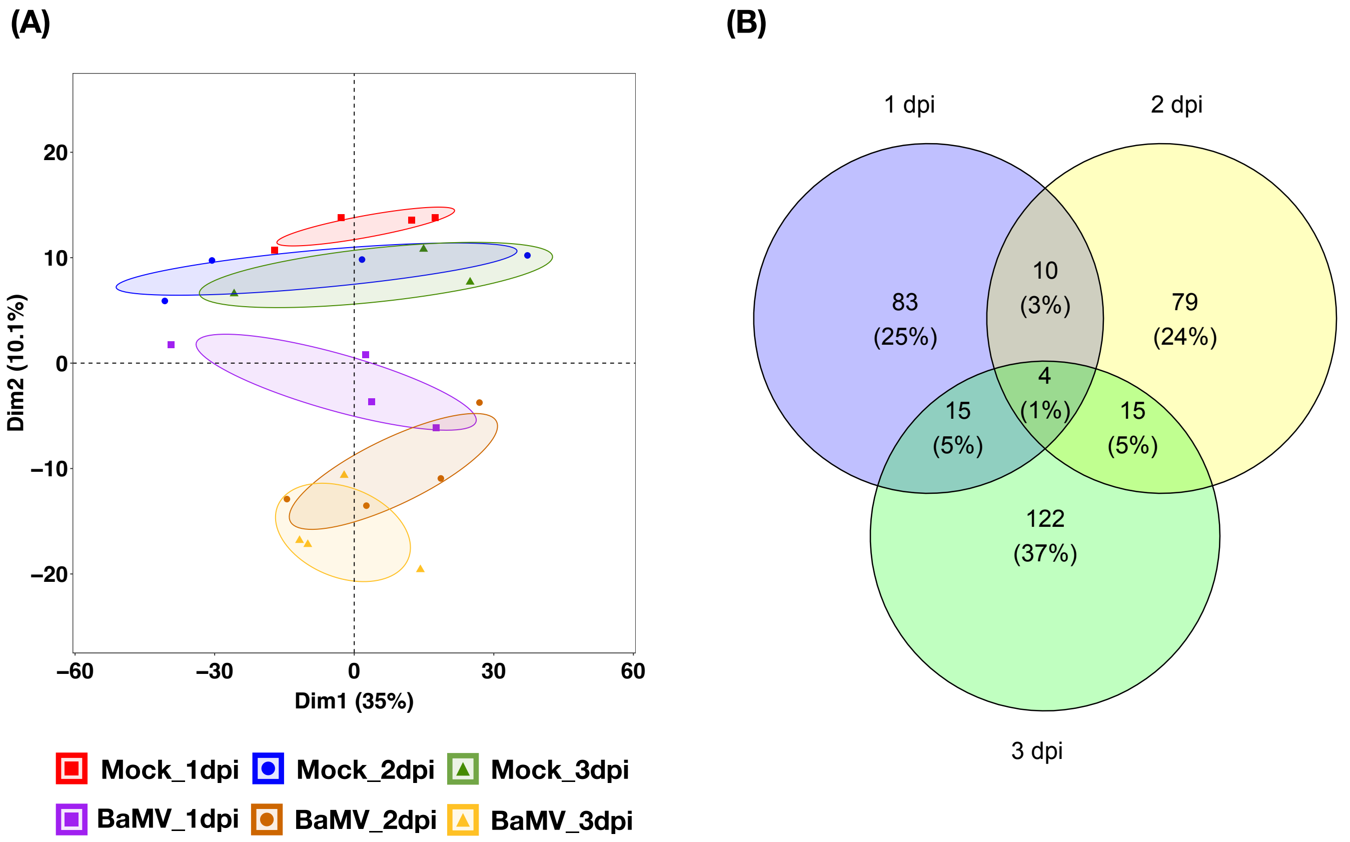


**Figure S1. The landscape of proteome between mock- and BaMV-infected leaves.** Leaves of *N. benthanmiana* plants were inoculated with 1 µg BaMV virions and harvested at 1, 2, and 3 day-post-inoculation (dpi). Leaf samples were used for protein extraction followed by a proteomic analyse. (A) The results of protein abundance were further subjected to principal component analysis. (B) The relationship of differential expressed proteins between BaMV-infected leaves harvested at different time points were illustrated using Venn diagram. Results are from 3 to 4 biological replicates.

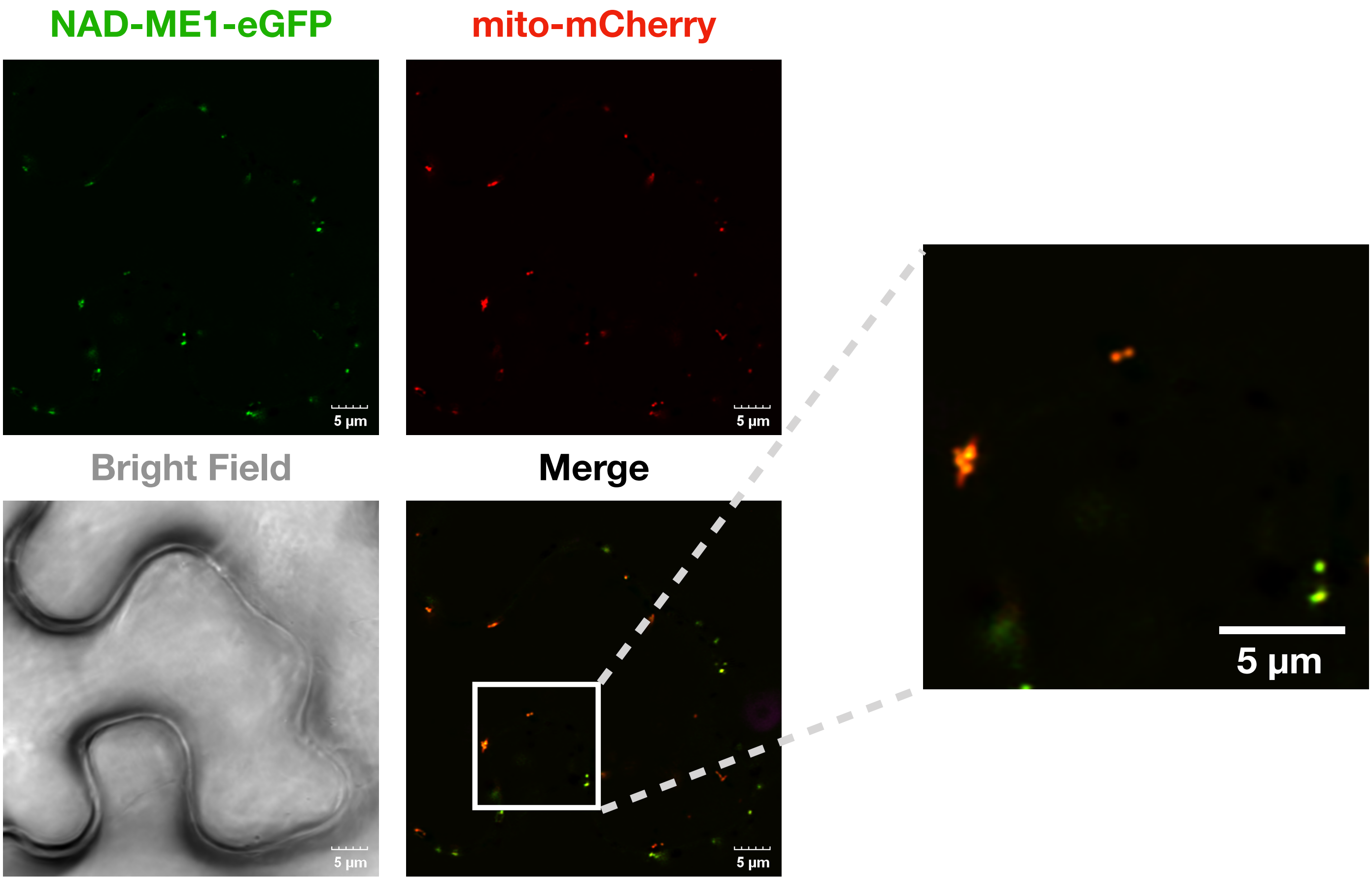


**Figure S2. The subcellular localization of NAD-malic enzymes 1 (NAD-ME1).** Leaves of *N. benthanmiana* plants were infiltrated with agrobacteria harboring clones of NAD-ME1-eGFP and mito-mCherry. At 2-day-post-infiltration, leaves were subjected to confocal microscopy.

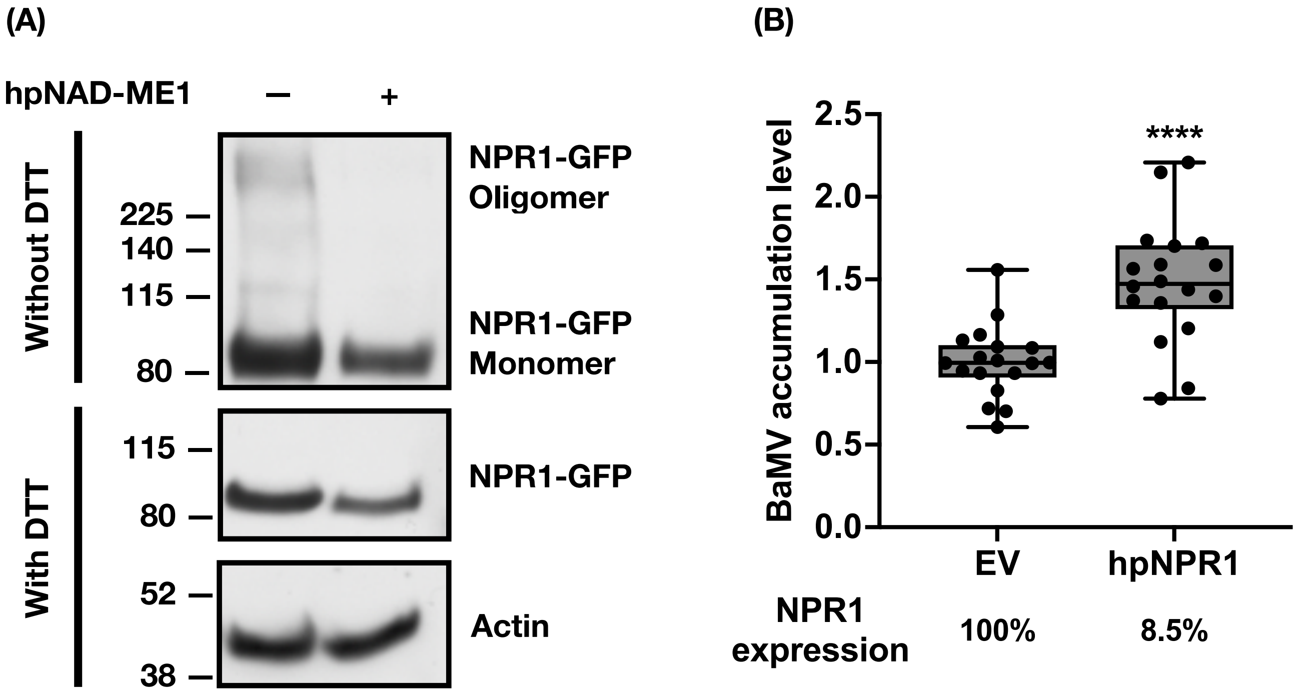


**Figure S3. The impact of silencing NAD-malic enzymes 1 (NAD-ME1) on protein stability of nonexpressor of pathogenesis-related genes 1 (NPR1) and the impact of silencing NPR1 on BaMV accumulation.** Leaves of *N. benthanmiana* plants were infiltrated with agrobacteria harboring hairpin RNA silencing clones. At 3-day-post-infiltration, leaves were inoculated with 1 μg BaMV virions and harvested at 27-hour-post-inoculation. (A) NPR1 protein accumulation in non-reducing (without DTT) and reducing (with DTT) conditions. Actin protein abundance serves as loading control. (B) BaMV accumulation in NPR1 silencing plants. Results are from 18 biological replicates. Statistical analysis was conducted using unpaired t test (*$P < 0.05$; **$P < 0.01$; ***$P < 0.001$; ****$P < 0.0001$).
