## Supplemental Table S2 for "Virus-induced upregulation of mitochondrial metabolism modulates cytosolic redox balance and defense responses"

**Supplementary Table S2.** The levels of starch and soluble sugars in leaves of mock- and BaMV-infected *N. benthamiana* plants. Results are the mean with SE from four biological replicates. Statistical analysis was conducted using unpaired t test (*$P < 0.05$, compared to mock-infected leaf). FW, fresh weight.

| Metabolites | Mock | BaMV |
|---|---|---|
| Starch (μmol C6 g $FW^{-1}$) | 450.9 ± 7.151 | 460.1 ± 7.877 |
| Glucose (μmol g $FW^{-1}$) | 0.306 ± 0.011 | **0.590 ± 0.008*** |
| Maltose (μmol g $FW^{-1}$) | 2.930 ± 0.040 | **3.213 ± 0.070*** |
| Fructose (μmol g $FW^{-1}$) | 0.358 ± 0.015 | 0.335 ± 0.017 |
| Sucrose (μmol g $FW^{-1}$) | 1.680 ± 0.033 | **1.819 ± 0.036*** |
